# A cross-species framework for investigating perceptual evidence accumulation

**DOI:** 10.1101/2024.04.17.589945

**Authors:** Sucheta Chakravarty, Cristina Delgado-Sallent, Gary A. Kane, Hongjie Xia, Quan H. Do, Ryan A. Senne, Benjamin B. Scott

## Abstract

Cross-species studies are important for a comprehensive understanding of brain functions. However, direct quantitative comparison of behaviors across species presents a significant challenge. To enable such comparisons in perceptual decision-making, we developed a synchronized evidence accumulation task for human and non-human animals, by aligning mechanics, stimuli, and training. The task was readily learned by rats, mice and humans, with each species exhibiting qualitatively similar performance. Quantitative model comparison revealed that all three species employed an evidence accumulation strategy, but differed in speed, accuracy, and key decision parameters. Human performance prioritized accuracy, whereas rodent performance was limited by internal time-pressure. Rats optimized reward rate, while mice appeared to switch between evidence accumulation and other strategies trial-to-trial. Together, these results reveal striking similarities and species-specific priorities in decision-making. Furthermore, the synchronized behavioral framework we present may facilitate future studies involving cross-species comparisons, such as evaluating the face validity of animal models of neuropsychiatric disorders.

**Highlights:**

1. Development of an evidence accumulation task for rats and mice
2. Synchronized video game allows direct comparisons with humans
3. Rat, mouse and human behavior are well fit by the same decision models
4. Model parameters reveal species-specific priorities in accumulation strategy

## Introduction

Quantitative comparisons of behavior across species are crucial to understand complex cognitive processes (Badre et al., 2015; Esteves et al., 2021; Schmack et al., 2021; Young, 2023). However, such comparisons are challenging for several reasons (Barron et al., 2020; Redish et al., 2021). Research in individual laboratories commonly focuses on single species, leading to marked differences in experimental techniques across species (Badre et al., 2015). These experiments often exploit the innate abilities of individual species. For example, humans are commonly provided with verbal task instructions, which is not possible with other animals. The disconnect between human and animal research can be costly, since animal models serve an important role in preclinical research of human neuropsychiatric disorders (Markou et al., 2009; Nestler & Hyman, 2010).

One domain where significant progress in cross species analysis has been made is perceptual decision making (Brunton et al., 2013; Nguyen & Reinagel, 2022; Pedrosa et al., 2023; Schmack et al., 2021; Shevinsky & Reinagel, 2019). Perceptual decision making is a core behavior observed in many species, and alterations have been observed across a range of human neuropsychological disorders (Horga & Abi-Dargham, 2019; Kavcic et al., 2011; Rizzo & Nawrot, 1998; Robertson & Baron-Cohen, 2017; van den Boogert et al., 2022). Previous research has investigated perceptual decisions in humans, non-human primates, and rodents, using similar stimuli (Carandini & Churchland, 2013; Hanks & Summerfield, 2017; Shadlen & Kiani, 2013). However, direct comparisons of cross-species behavior based on previous studies is difficult, due to marked differences in the paradigms and the training protocols.

Here we present a behavioral framework for investigating perceptual decision making in mice, rats and humans. We implemented a free response version of the pulse-based evidence accumulation task, previously used in rodents, especially in rats (Brunton et al., 2013; Gupta et al., 2024; Scott et al., 2015; Kane et al. 2023) and mice (Odoemene et al., 2018; Pinto et al., 2018). Briefly, the task uses sequences of brief visual pulses (flashes), presented to the left and right sides while the subject must choose the side with the higher pulse probability to obtain the reward. This task offers a speed-accuracy trade off as longer response times allow the observer to sample more pulses, which enables better estimates of the relative pulse probability. We trained rats and mice to perform this task using computer controlled 3-port light chambers, and an automated training facility. In parallel, we developed a video game based on the same task to gather data from human participants online. We synchronized task parameters and training protocols between the rodent task and the human video game; we used non-verbal, reward feedback-driven training in all three species. Further, the high-throughput rodent training facility, and the online game facilitated large-scale data collection.

All three species learned to perform the task and relied on evidence accumulation as a choice strategy. However, there were cross-species differences in several key aspects of the accumulation process. Overall, humans were slower and more accurate than rodents, particularly mice. Drift diffusion models (DDM) revealed highest decision thresholds in humans, and lowest thresholds in mice. Further, collapsing boundary model fits indicated that rodent behavior may be limited by internal time-pressures. Interestingly, rats appeared to optimize response times to maximize reward rate, whereas humans opted for better accuracy. Mice showed high animal-to-animal and trial-to-trial variability in choice behavior and model fits suggested they alternated between evidence accumulation- and other strategies across trials. Together these data reveal important similarities and species-specific priorities in decision making across rodents and humans, and highlight the value of synchronized behavioral frameworks for cross-species studies.

## Results

### A synchronized framework for cross-species studies of perceptual decision making

We developed a free-response version of the pulse-based evidence accumulation task (Kane et al. 2023) that was synchronized across rodents and humans (Figure 1; see methods for details). In this task, sensory information was presented as a sequence of randomly-timed pulses of light from two sources, one to the left and one to the right (Figure 1B). Pulses were brief (10 *ms*) and binned into 100 *ms* bins—the odds were set so that the probability (*p*) of a flash on one side in a bin was complementary to the flash probability on the other side for the same bin (1 − *p*). The pulses continued until the subject selected one of the light sources. Identification of the light source with the greater probability of pulsing yielded a correct response.

**Figure 1:**
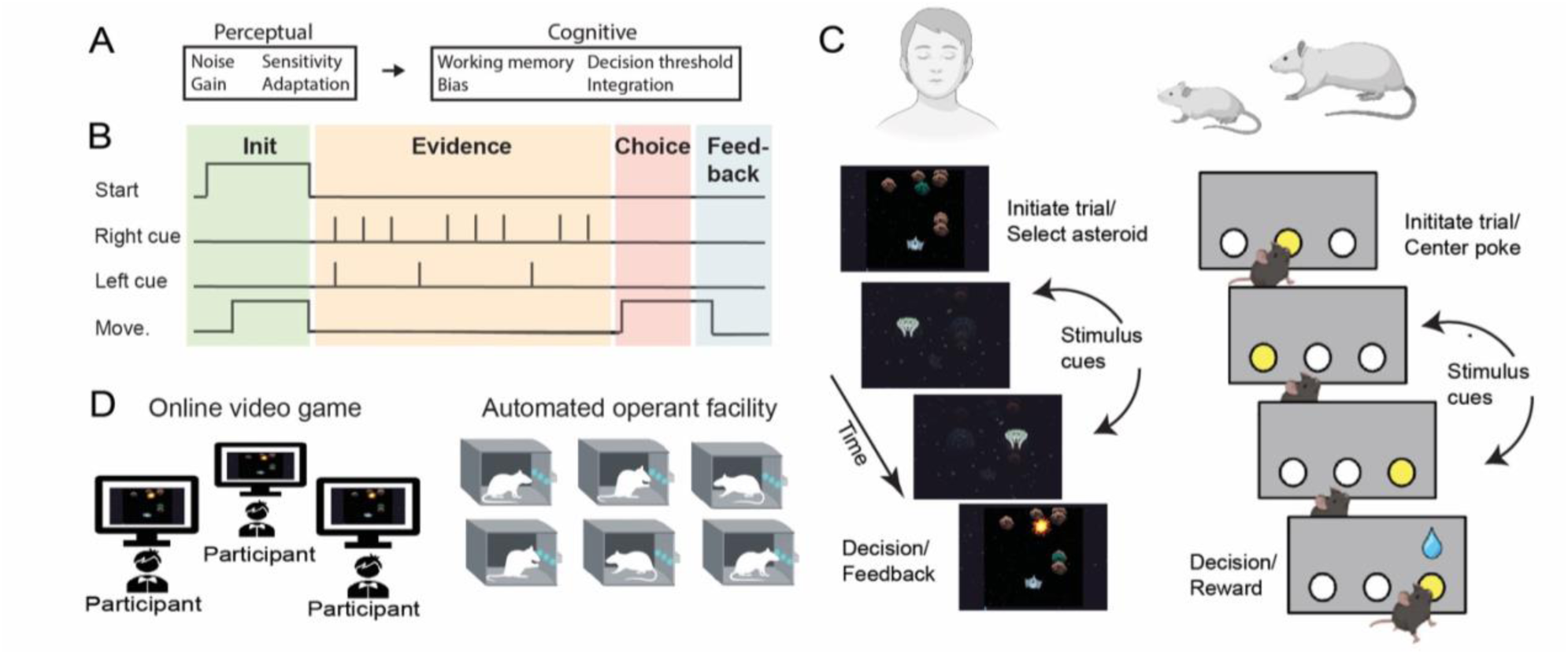
Synchronized pulse-based evidence accumulation task for humans and rodents. A) During perceptual decision-making, choices are influenced by perceptual factors, for example sensory noise and sensitivity, as well as cognitive factors, including decision thresholds and working memory. Pulse based evidence accumulation tasks were developed to discriminate the relative contributions of perceptual and cognitive factors on behavioral choice across individuals. B) Schematic of the pulse-based evidence accumulation task for cross-species studies. The goal of the task is to identify the side with the higher probability of visual stimuli. The task structure is the same across species and consists of 4 different phases: *Initiation*: the subject starts the task by clicking an asteroid (humans) or poking in the center (rodents), *Evidence*: the visual stimuli will be presented in the right or left side. *Choice*: during the evidence period, the subject observes the cues and responds by selecting a side. Cues continue until the side is selected. The correct choice is the side with the greater underlying probability of cue presentation. *Feedback*: the subject will receive a reward (increase of the score for humans and sugar water for the rodents) for a correct choice or a time-out (TO) delay for an incorrect choice. C) *Left:* The human version consists of an online video game format in which the player controls a spaceship in an asteroid field. The player gets points by destroying asteroids with a laser beam. *Right:* In the rodent version, mice and rats play the task in a 3-port operant training chamber, in which the center port is used for initiation and the left and right ports are used for making choices. D) *Left:* For humans, data was collected using remote video calls format. *Right*: For rodents, behavioral sessions were conducted from multiple animals in parallel in a semi-automated operant facility.

Rodents performed the task in a three port operant chamber; they initiated each trial with a nose poke at the center port, followed by a cue period during which sequences of brief light flashes were presented in the left and right light ports, the flashes continued until the animal poked on one of the side ports. Rodents were rewarded with a drop of sugar water for poking on the side with greater flash probability (Figure 1C). For the human experiments, we designed an online video game that preserved the same mechanics and stimulus statistics (flash duration, flash rate, and generative flash probability) from the rodent task (Figure 1C). In the video game, participants were presented with a field of asteroids, each trial started when they mouse-clicked on any asteroid, following which sequences of brief flashes in the form of an alien spaceship were presented bilaterally—this was the cue period; the flashes continued until the participant mouse-clicked on one of the sides. If they chose the side with the greater flash probability, the initially selected asteroid would be destroyed (Figure 1C). Importantly, both the rodent task and the human video game offered a speed-accuracy tradeoff. Longer response times facilitated greater probability of choosing the correct side, due to greater opportunity for accumulating evidence from the flashes.

Both the rodent task and the human video game used a non-verbal, feedback-based training pipeline (Do et al., 2023; Kane et al. 2023). This pipeline consisted of progressive phases to familiarize subjects to task mechanics and game rules (Supplementary Figure 1). Correct choices were rewarded with positive feedback—a drop of sugar water for the rodents and point bonuses for the humans. Importantly, humans did not receive verbal instructions about the game rule. Rodents underwent multiple sessions of training over the course of many days (4 − 5 weeks for mice and 1 − 3 weeks for rats), while humans completed 1-2 sessions (several minutes).

### Rodents and humans solved the task using an evidence accumulation strategy

We trained 21 rats, 95 mice and 18 adolescent humans on the task. All three species performed well during the testing phase and were above the criteria for chance performance (upper bound of the 99% binomial confidence interval, see methods). There was substantial heterogeneity in performance across individuals of a species, revealed by their overall accuracy, response time (RT), bias and reward rate (Figure 2A-D). On average, humans were slower and more accurate, mice were the fastest and least accurate (One-way ANOVA for accuracy: *p* < 0.0005, *F* = 271.84; for RT: *p* < 0.0005, *F* = 236.39, see Figure 2A-B, E). Interestingly, there was overlap in performance across species (e.g. Figure 2A-2C). There were no detectable differences between males and females in any species (Supplementary Figure 2).

**Figure 2:**
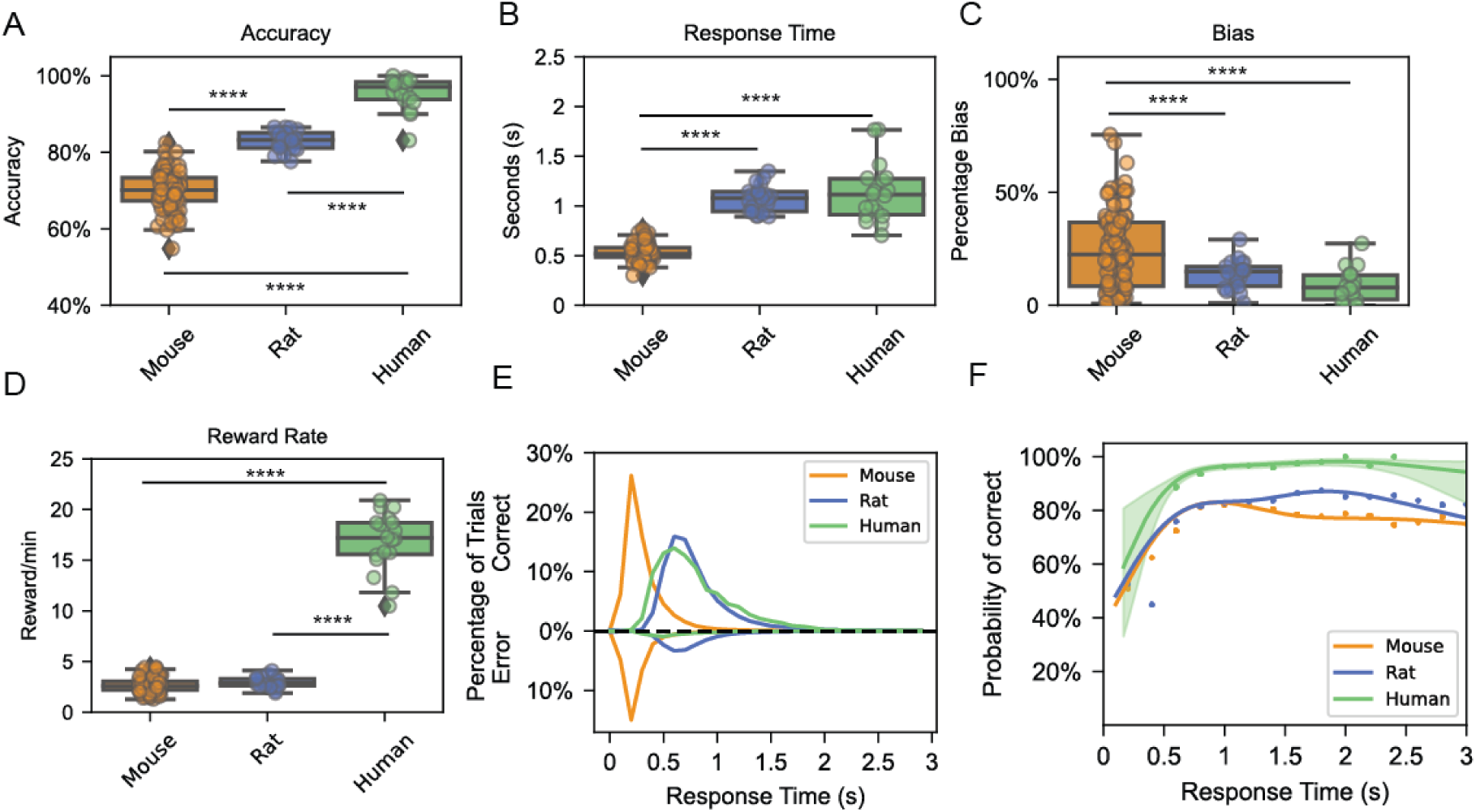
Humans, mice and rats accumulate evidence over time to solve the task. A) Accuracy varies significantly across species (One-way ANOVA; p < 0.0005, F = 271.84), with mice exhibiting the lowest performance and humans demonstrating the highest. B) Mice exhibit faster response times compared to humans and rats, which display similar response times (One-way ANOVA; p < 0.0005, F = 236.39). C) Normalized bias is comparable between humans and rats but is higher in mice (One-way ANOVA; p < 0.0005, F = 11.13). D) Reward rate was similar for rodents but larger for humans (One-way ANOVA; p < 0.0005, F = 1060.97). E) Histogram of response times across species reveals that rats and humans respond at similar speeds during correct or incorrect trials, whereas mice respond faster in error trials than in correct trials (Paired t-student; p < 0.0005). Note that positive values represent correct trials, while negative values represent incorrect trials. F) Accuracy vs response time derived from a generalized additive mixed model on the three species in which we can observe the evidence accumulation process in each of the species. All of the species peak between 0.7-1 seconds but the humans present a higher peak than the rodents. Each dot represents the averaged accuracy per each 0.2-second bin for each species (Pearson Correlations Accuracy-RT, see methods; Mouse: *p* < 0.017, *R* = 0.80, Rat: *p* < 0.011, *R* = 0.83, Human: *p* < 0.033, *R* = 0.85). All comparisons are unpaired two-sample t-tests, corrected for multiple comparisons with Bonferroni correction. Notations: * = p<0.05, ** = p<0.01, *** = p<0.005, **** = p<0.001.

Importantly, two pieces of evidence suggested that all three species used a similar evidence accumulation strategy. First, across species, trials with longer RTs yielded increased accuracy (Pearson correlations between accuracy and RT, see methods; Mouse: *p* < 0.017, *R* = 0.80, Rat: *p* < 0.011, *R* = 0.83, Human: *p* < 0.033, *R* = 0.85). This was illustrated with a generalized additive mixture model (GAMM), fit to the choice and RT data (Figure 2F). Second, choice and RT data from individuals of each species were well fit by the drift diffusion model (DDM; Bogacz et al., 2006; Ratcliff, 1978; Ratcliff & McKoon, 2008), which showed relatively high decision bounds (Figure 3A-C).

**Figure 3:**
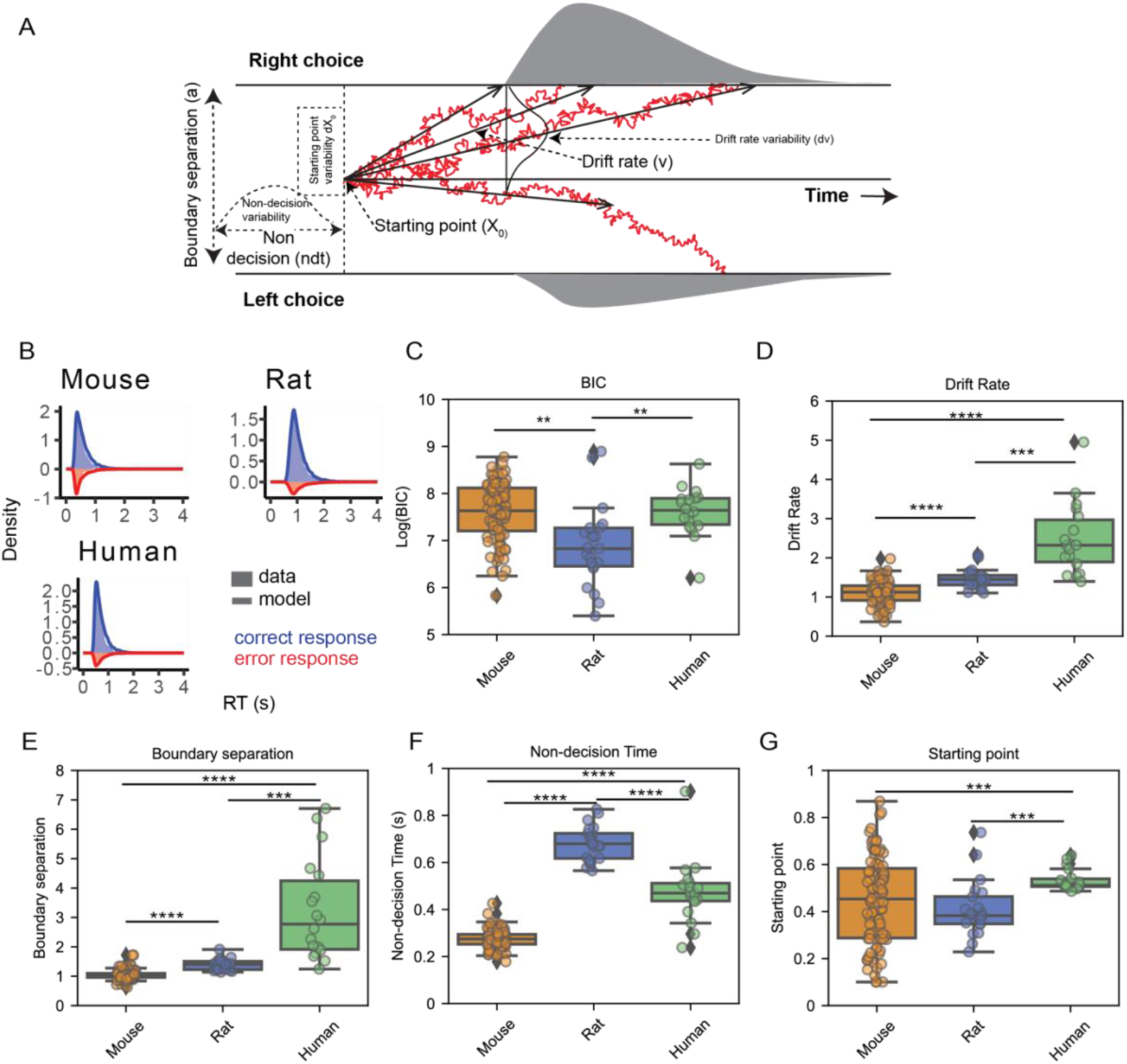
Drift diffusion models explain the choice and response time data for all three species. A) Illustration of the diffusion model: two boundaries indicate decision thresholds for left and right choice, greater distance between boundaries would mean more separability between the two choice decisions. Drift rate is the average rate of evidence accumulation, the model assumes drift rate varies from trial to trial, following a normal distribution. Starting point measures the bias in choosing one side over the other. Starting point is also assumed to vary from trial to trial, following a uniform distribution. Non-decision time indexes time spent in processes other than the decision process. Trial to trial variability in non-decision time follows a half-normal distribution. Evidence on each trial is accumulated until a decision boundary is reached, evidence accumulation is governed by non-decision time, starting point bias, drift rate; drift across time is subject to a diffusion noise. B) Fit of the diffusion model to one example subject from each species, response time distributions are plotted separately for correct and error responses. C) Comparison of model performance across species. Rats are better fitted by the model than mice and humans (DDM BIC; Humans: 2772 ± 262, Rats: 1492 ± 403, Mice: 2778 ± 136; One-way ANOVA; p < 0.0005, F = 10.97). D-G) Estimates of model parameters compared across species. Rodents present lower drift (One-way ANOVA; p < 0.0005, F = 77.72) and boundaries (One-way ANOVA; p < 0.0005, F = 88.29) than humans. Rats show the slower non-decision time (One-way ANOVA; p < 0.0005, F = 347.26). All comparisons are unpaired two-sample t-tests, corrected for multiple comparisons with Bonferroni correction. Notations: * = p<0.05, ** = p<0.01, *** = p<0.005, **** = p<0.001.

### Species differ in key model parameters revealing differences in the accumulation process

While all three species were well fit by the DDM, our initial investigation showed cross-species differences in RT, suggesting possible differences in the accumulation process across species. To further explore cross-species differences, we examined how DDM parameters varied across species. The DDM assumes that the accumulation process is governed by four main parameters: **boundary separation** (*a*)**, drift rate** (*v*), **non-decision time** (*ndt*), and **starting point bias** (*x*_0_); these were estimated for individual subjects (Figure 3A). Comparison of the DDM parameters showed that humans were characterized by the largest drift rate (*v*) and boundary separation (*a*), whereas mice had the lowest drift rate and boundary separation (Drift rate: *p* < 0.0005, *F* = 77.72; Boundary separation: *p* < 0.0005, *F* = 88.29; Figure 3D-E). Rats showed the greatest non-decision time (*ndt*), while mice had the shortest non-decision time (*p* < 0.0005, *F* = 347.26; Figure 3F). There was also a trend effect for greater starting point bias (*x*_0_) in rodents (*p* = 0.053, *F* = 2.99; Figure 3G).

Thus, the DDM parameters suggested key differences in the accumulation process across species. The higher boundary separation in humans suggested a higher threshold for perceptual evidence before commitment to a decision, leading to more accurate but slower responses. The greater non-decision time and lesser boundary separation in rats suggested that rats accumulated less evidence, but spent more time in processes unrelated to decision making (for example, motor movements). Finally, mice had the least boundary separation indicating less evidence accumulation, also less time spent in non-decision processes. Accordingly, mice were the fastest and least accurate of the three species.

### Rodents’ choices were limited by time-pressures

Given the faster RT in mice, and model-based evidence for faster decision times in rats, we asked if rodents’ choices were limited by internal time-pressures. The classic version of the DDM discussed above, does not readily account for this possibility, for it assumes that with time in a trial, the amount of evidence required to make a decision remains the same. This assumption is implemented in the model through a fixed decision boundary (*a*). To test if rodents experienced time-pressures, we considered two additional variants of the DDM (see methods and Supplementary Figures 2, 3 and 4), a collapsing boundary model (Bowman et al., 2012; Shevinsky & Reinagel, 2019), and an urgency model (Ditterich, 2006).

The collapsing boundary DDM assumes that with time in a trial, the decision boundary collapses, and thus, less evidence is required to make a decision. It included three additional parameters: **initial boundary separation** (*a*_0_), **semi-saturation constant** (*t*_0.5_), and **amount of collapse** (*k*) of the boundary (see methods and Figure 4A). The urgency model incorporates an urgency signal that increases with time, raising the probability of a choice. The collapsing boundary- and urgency models are mathematically similar, but the collapsing boundary model can better explain faster RTs on more difficult trials. The urgency DDM also included three additional parameters: **slope** (*u*_*slope*_), **magnitude** (*u*_*mag*_), and **onset** (*τ*) of the urgency signal (Figure 4B).

**Figure 4:**
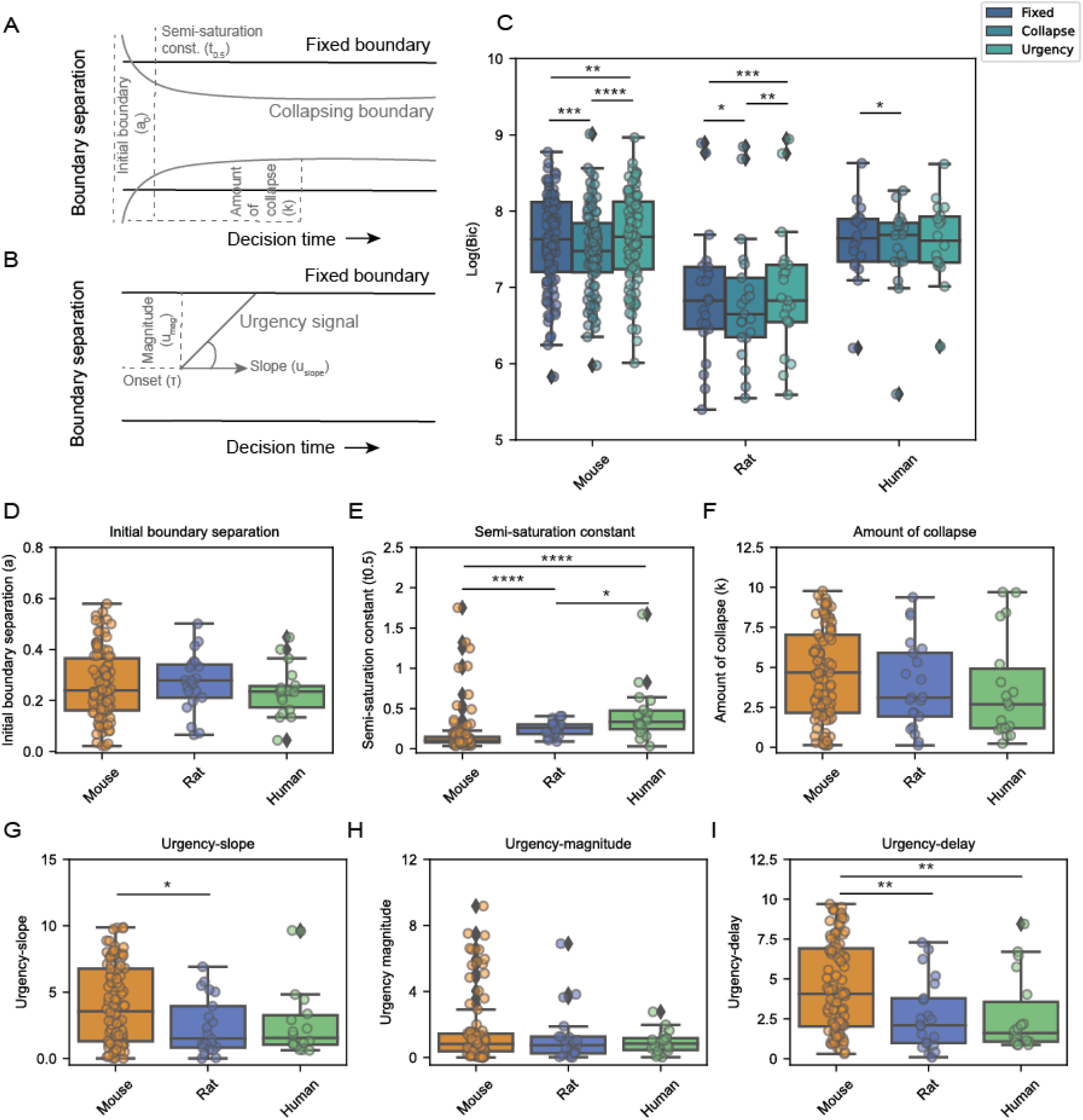
Response times for rats and mice are well described by collapsing bounds. A) Illustration of the diffusion model with collapsing boundary, three (free) parameters are used to estimate the boundary: initial boundary separation, semi-saturation constant, and amount of collapse; the model is otherwise the same as the fixed boundary model. B) Illustration of the diffusion model with an urgency signal, three (free) parameters are used to estimate the (linear) urgency signal: slope of the signal, magnitude of the signal, and onset delay for the signal. C) Comparison of model performance for the three diffusion models: fixed boundary model, collapsing boundary model, and the urgency model, for each species. Rats and mice were better fitted by the collapsing boundaries (Repeated measures ANOVA; Mice: p < 0.0005, F = 22.12; Rats: p = 0.0014, F = 11.12). D-F) Comparison of boundary related parameters of the collapsing boundary model, across species. Mice show the lowest semi-saturation constant (One-way ANOVA; p = 0.004, F = 5.69). G-I) Comparison of urgency related parameters of the urgency model across species. Mice present the highest urgency-slope (One-way ANOVA; p = 0.027, F = 3.71) and urgency delay (One-way ANOVA; p = 0.003, F = 6.06). Notations: * = p<0.05, ** = p<0.01, *** = p<0.005, **** = p<0.001.

Model comparison using the BIC revealed that the collapsing boundary DDM fit rodent data significantly better than the classic- or the urgency DDM (Repeated measures ANOVA; Mice: *p* < 0.0005, *F* = 22.12; Rats: *p* = 0.0014, *F* = 11.12; Figure 4C). However, for humans, the collapsing boundary- or the urgency model was not better than the classic DDM (One-way ANOVA; *p* = 0.13, *F* = 2.31; Figure 4C). These results suggest that rodents’ behavior was limited by internal time-pressures. Further, the collapsing boundary DDM showed the lowest semi-saturation constant (*t*_0.5_) for mice, which suggests that the boundary collapse was fastest for mice (*p* = 0.004, *F* = 5.69; Figure 4E), producing the fastest responses.

Overall, these results supported the hypothesis that internal time-pressures limited rodents’ performance in the current task.

### Rats balance speed and accuracy near optimally to maximize the reward rate

Summary performance measures suggested cross-species differences in accuracy and RT, and the DDMs helped explain the differences in the underlying decision parameters. However, it remains unclear how each species settled on their particular speed-accuracy tradeoffs (Figure 5A). One possibility is that they chose an RT that maximized their reward rate (Figure 5B-C and Supplementary Figure 6, see also Bogacz et al. 2006).

**Figure 5:**
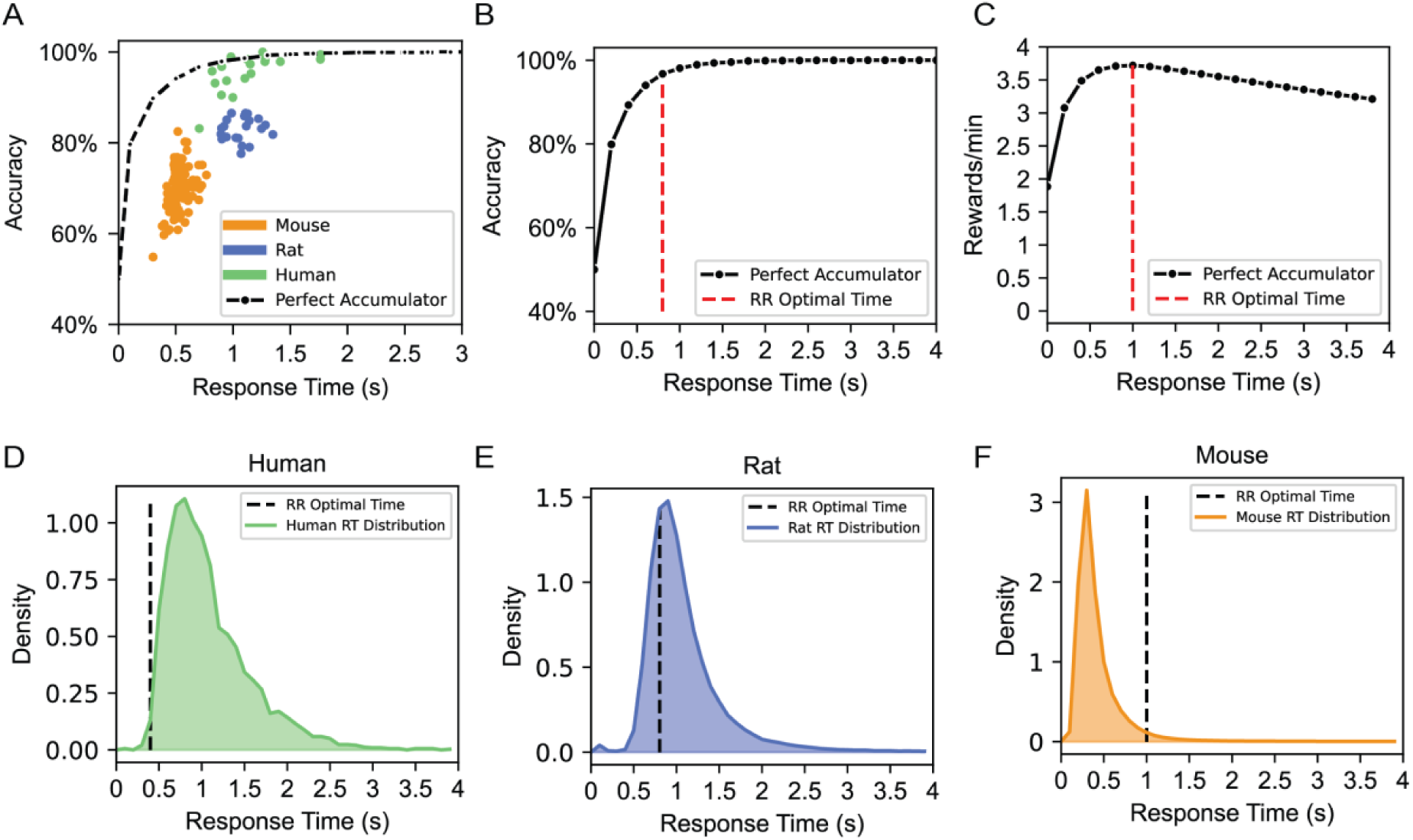
Rats balance speed and accuracy near optimally to maximize reward rate. A) The speed-accuracy trade-off plot illustrates the relationship between response time and accuracy across species, each dot represents the mean accuracy and response time of an individual. The dashed line represents the accuracy as a function of different choices of RT for a perfect accumulator. B-C) Representations of accuracy (B) and reward rate (C) as a function of different choice of RT for a perfect accumulator, incorporating the average trial initiation time observed in mice, which was different for correct and error responses. In the simulations, flash ratio was kept at 80% vs 20%, and flash rate was 10 Hz. The red line marks the RT associated with the peak reward rate of the accumulator. D-F) Histograms illustrate the distribution of RT observed in humans (D), mice (E), and rats (F) in comparison to the RT associated with the peak reward rate of the perfect accumulator (black dashed lines).

To investigate this possibility, we simulated a perfect accumulator for each species, accounting for cross-species differences in the inter-trial interval (Supplementary Figure 6G-I, also see methods). Interestingly rats were close to the simulated optimal RT, while humans responded slower than the optimal RT and mice were faster (Figure 5D-F). These observations suggested that rats may balance speed and accuracy to maximize reward rate.

### Mice were more influenced by alternate strategies across latent states

Previous research suggests existence of alternate decision strategies in evidence accumulation tasks (Ashwood et al., 2020; Carandini & Churchland, 2013; Roy et al., 2021). To investigate if subjects in our task displayed shifts in strategy, we used a generalized linear model (GLM) that estimated the binomial probability of choosing the right-over the left side, as a linear function of perceptual evidence and the following alternate strategies: **win-stay-lose-switch** (determined by the choice and reward), and **previous choice** (determined by choice independent of reward). Perceptual evidence was the flash difference, divided by total flashes 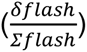. The intercept term represented side-bias (Figure 6A, also see methods). The GLM suggested that rats and humans relied more on perceptual evidence, but mice were influenced by alternate strategies (*p* < 0.0005, *F* = 156.28; Figure 6D-E). The GLM predicted the psychometric curve for rats and humans well, but for mice the fit was poorer (*p* < 0.0005, *F* = 38.67; Figure 6B-C).

**Figure 6:**
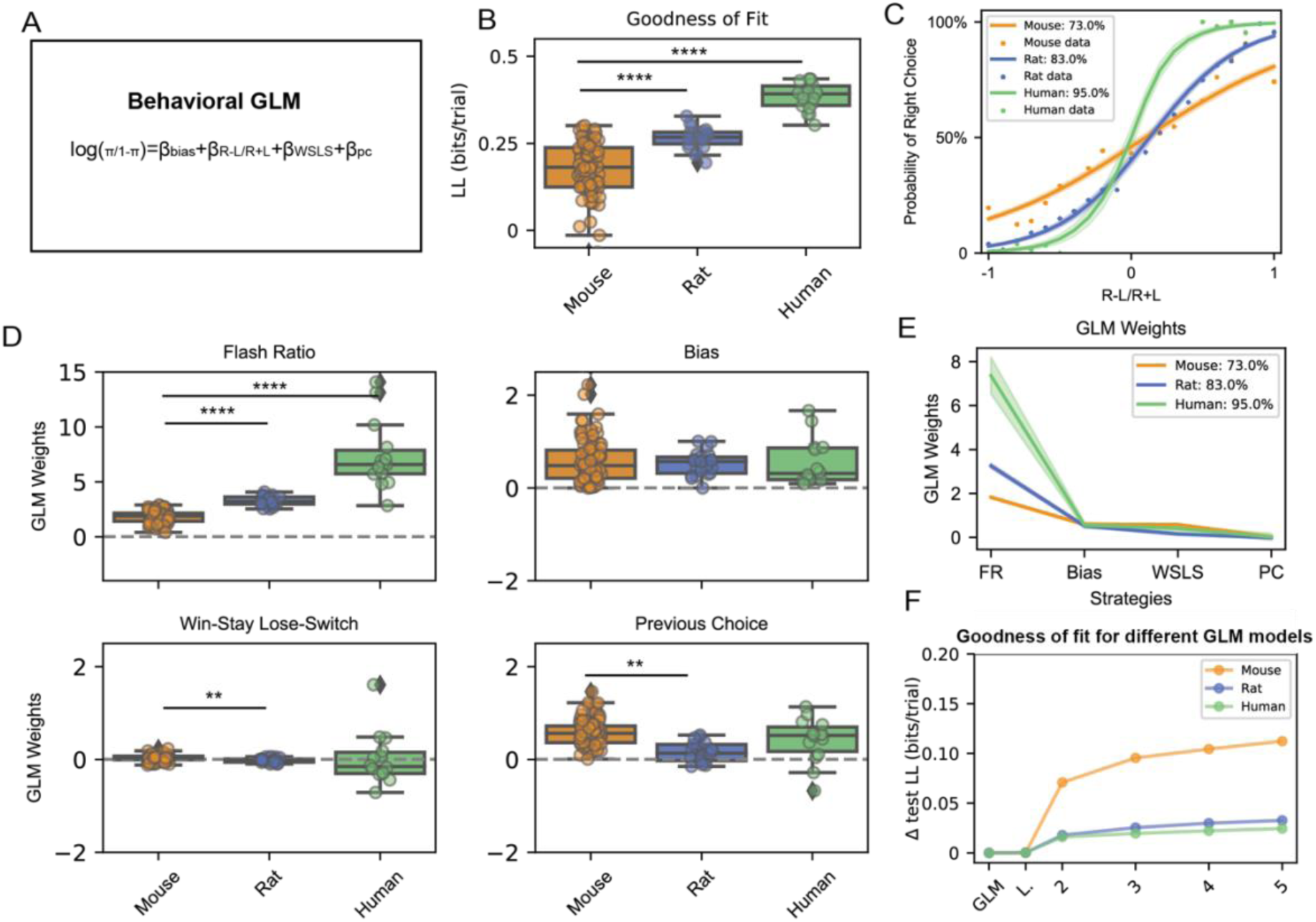
Mice alternate between different strategies while performing the task. A) Generalized linear model with 4 parameters: Bias (Intercept), Flash Ratio (Right flashes - Left Flashes) / Total flashes), Win-stay Lose-switch, and Previous choice. B) Goodness of fit of the GLM presents a better fit for the human data compared to mice or rats. Test log-likelihood is in units of bits per trial, and is relative to a ‘null’ Bernoulli coin-flip model (One-way ANOVA; p < 0.0005, F = 38.67). C) Psychometric curve fits, and mean predictive accuracy across species. D) Estimated GLM weights showed that humans relied on the flash ratio more than the rodents, followed by the rats (One-way ANOVA; p < 0.0005, F = 156.28), whereas mice tended to use a previous choice strategy (One-way ANOVA; p < 0.0005, F = 18.2). E) Mean estimated GLM weights across species. F) Model comparisons among the one-state GLM, the classic lapse model (’L’), and multiple GLM-HMMs with different numbers of latent states. To compare across species, we measured the change in test log-likelihood, as a function of the number of states included in the model, relative to a one-state GLM, separately for each species. Mice showed greater increase in LL for a GLM-HMM with 3 latent states, suggesting that mice transitioned through multiple latent states while performing the task. All comparisons are based on unpaired two-sample t-tests, corrected for multiple comparisons with Bonferroni correction. Notations: * = p<0.05, ** = p<0.01, *** = p<0.005, **** = p<0.001.

The influence of multiple strategies on decisions may suggest that subjects dynamically switch behavioral strategies on a trial by trial basis (Ashwood et al., 2022). To test this hypothesis, we used a modeling framework previously proposed by Ashwood et al., (2022) that combined a hidden Markov model (HMM) with a GLM. Using this GLM-HMM, we investigated the number of latent states underlying choice behavior in each species. We considered the original GLM, a GLM with lapse states, and multiple GLM-HMMs with increasing latent states—2, 3, 4, and 5 states, respectively. Model comparison showed that the original GLM fit the data from rats and humans better (Figure 6F). However, mice data were better fit by a 3-state GLM-HMM (Figure 6F); the 3 states were evidence accumulation, and left- and right side biases (Supplementary Figure 7). Thus, mice appeared to alternate between different latent states that prioritized evidence accumulation or alternate decision strategies.

In line with the above results, we also found interesting, time-varying trends in raw behavior (Supplementary Figure 8). Rats were remarkably stable in maintaining a constant RT and accuracy across their 120-minute session. Rats performed similar numbers of trials across time and had a stable reward rate across the session. In contrast, in mice, RT, accuracy and reward rate changed over the course of a session. Thus, mouse behavior in this task was more variable trial-to-trial when compared to rats or humans.

## Discussion

We present a behavioral framework, synchronized across rodents and humans, for investigations of perceptual decision making. Inspired by pulse-based evidence accumulation tasks, previously used in rats, we designed a free response task that offered a speed-accuracy trade-off. In parallel, we developed a video game based on the same task to test humans online. Together we used this task and game to assess species-specific priorities in decision making. Task parameters and training procedures were deliberately synchronized across species, facilitating direct, quantitative comparisons of behavior. All three species learned to perform the task, their behavior was well fit by evidence accumulation models, and we found significant species-specific differences in key parameters. Unlike humans, rodents were best fit by a collapsing boundary DDM suggesting that their performance was limited by internal time-pressures. Rat’s response times were close to optimal to maximize reward rate. Mice alternated between evidence accumulation- and other strategies, and exhibited more variable behavior. Overall, this study highlights the importance of a synchronized behavioral framework for meaningful, quantitative comparisons of cross-species perceptual decision making.

Our framework has several key attributes to facilitate quantitative comparisons of behavior across species. First, to minimize training differences, we used a similar non-verbal feedback-based training strategy in which difficulty was slowly increased to facilitate learning. Second, in humans we used an online video game-based approach. Video games can be advantageous over traditional psychological experiments; games are engaging and can be used to collect objective behavioral measures with well-controlled stimuli. Also, online games are helpful for collecting larger datasets (see also, Do et al., 2023). Finally, the short, self-paced nature of the video game enabled data collection within a relatively brief period of time (mean completion time = 11.06 minutes +/- 2.26 S.D., while the task could be implemented in a high-throughput, semi-automated rodent training facility allowing collection of behavioral data from many animals. Together, this framework is advantageous for collecting larger datasets across different species within a relatively short amount of time.

An interesting result revealed by this approach is that rodent performance was best fit by a collapsing bound model, suggesting their decisions were limited by internal time-pressures. This was revealed by a comparison of the classic- (Ratcliff, 1978) and the collapsing boundary (Bowman et al., 2012; Shevinsky & Reinagel, 2019) version of the DDM. The classic DDM, a highly successful model for explaining perceptual decision making behavior (Ratcliff, 2014; Ratcliff et al., 1999; Ratcliff & Rouder, 1998), assumes that the amount of evidence required to make a decision remains constant. Others have suggested situations in which less evidence is required with time (Busemeyer & Rapoport, 1988; Drugowitsch et al., 2012; Thura et al., 2012, Ditterich, 2006). However, experimental evidence for the collapsing boundary model has been inconclusive (Hawkins et al., 2015; Voskuilen et al., 2016). Notably, while previous studies used these models to explain human and non-human primate data, here we show that rodents decisions are best fit by a collapsing bound model

While humans and rats were best fit by a single perceptual accumulation strategy across all trials, we found that mice were best fit by a model that alternated between evidence-accumulation and other strategies across different latent states. This finding is consistent with previous studies that have examined mouse behavior in other perceptual decision tasks (Ashwood et al., 2022; Carandini & Churchland, 2013; Gupta et al., 2024; Hanks & Summerfield, 2017).

The framework we provide here provides a complement to cross species studies using ethological approaches. Recent research has highlighted the importance of focusing on ethological behaviors in different species (Juavinett et al., 2018). Indeed, focus on ethological behaviors can provide unique insights into cross-species comparisons. However, this rarely allows for direct, quantitative cross-species comparisons as ethologically-relevant behaviors are often species-specific, reflecting adaptations to that organism’s particular niche. Here, we followed an approach that is complementary to the ethological focus, by designing a task that can be performed in similar ways by multiple species, in order to better appreciate the unique and common aspects of behavior.

### Limitations

Species specific differences we report here could reflect uncontrolled environmental variables rather than genetic differences. Despite a deliberate attempt to synchronize most aspects of our task and training across species, there were several differences in the human and rodent setups that could have contributed to the cross-species behavioral differences. First, rewards were different for rodents and humans; drops of sugar water for rodents vs point bonuses for humans. Moreover, while mice were water scheduled to increase motivation and rats were food scheduled. Second, rodents and humans performed the task for different durations, which could have impacted their motivational states differentially. Third, humans participated in the task from their home environment, with the experimenter only present remotely via Zoom. In contrast, the rodent underwent requisite animal handling procedures which were conducted by trained researchers.

### Conclusion

In sum, we present a synchronized behavioral framework for high-throughput data, and direct, quantitative comparisons of perceptual decision-making across species. This pulse-based free response evidence accumulation task is designed to differentiate between perceptual (sensory noise) and cognitive components (decision thresholds), which are distinct yet interacting processes affected in neuropsychiatric conditions such as autism spectrum disorder (Robertson & Baron-Cohen, 2017), Schizophrenia (Horga & Abi-Dargham, 2019), and Alzheimer’s disease (Kavcic et al., 2011; Rizzo & Nawrot, 1998). For example, in autism, perceptual symptoms are characterized by sensory hypersensitivity and hyposensitivity, altered sensory integration, and cognitive difficulties in filtering information, each of which can contribute to challenges in adapting to dynamic environments (Green et al., 2015; Keehn et al., 2013; Pellicano & Burr, 2012). The non-verbal, feedback-driven training pipeline for humans may allow inclusion of participants with communication difficulties, such as non-verbal autistic individuals, improving the reach of this research to underrepresented groups (McKinney et al., 2021). Moreover, such a framework not only supports basic neuroscience inquiries but also holds translational potential, enabling the quantification of behaviorally relevant traits that serve as objective benchmarks or biomarkers, to aid in better selection and assessment of genetic animal models. Finally, as this task is amenable to neurophysiological investigations, circuit mechanisms of behavioral traits can be followed up in the preclinical animal model.

## Acknowledgements

We thank Christa Rose, Arielle Rubel, Julie Gomez, Sanaa Ahmed, Anosha Khawaja-Lopez, Benjamin Lee and Sina Analoui for help with collection of rodent data, and Vanessa Torres-Lacarra and Phoebe Lesk for collection of human data. We thank Helen Tager-Flusberg for feedback on human data collection, Joseph McGuire for feedback on the analyses, and Josh Sanders for feedback on the behavioral training software. This work was supported by NIMH award number R56MH132732 and SFARI award number 874568 to BBS and a Center for Systems Neuroscience distinguished fellowship award to CDS and GAK.

## Contributions

SC, CDS, GAK, HX and QHD designed the experiments. SC and CDS analyzed the data. SC, CDS and BBS wrote the manuscript. All authors contributed to the conceptualization and execution of the study and edited the manuscript.

## Materials and methods

### Rodents

All experiments and procedures were performed in accordance with protocols approved by the Boston University Animal Care and Use Committee. Long Evans adult rats (N=21; aged 3 months to 2 years) were purchased from Taconic or bred in house. C57BL/6NJ adult mice (N=95; aged 2 to 8 months) were purchased from Jackson Laboratory. Both male and female rats or mice were utilized and trained simultaneously in the same room, albeit in separate operant chambers. Rats were food restricted to 80-100% of their body weight and fed once per day (typically 3-4 pellets of food per day). Mice were water restricted to 80-90% of their body weight and received a minimum of 1mL of water a day. Rats received 0.03 mL reward, whereas mice received 0.005 mL reward (10% sucrose solution; 100g/L in water). Reward volume was consistent within a session. Both rats and mice were housed on a 14:10 ON:OFF light schedule with the ON phase corresponding to daylight hours in Boston, MA USA.

### Rodent behavioral control system

Behavioral control system was inspired by Dhawale et al., 2017 and Poddar et al., 2013. Rodents were trained in custom acrylic chambers with three nose ports. Nose ports were 3D printed (Sanworks or custom made) and equipped with a visible LED for stimulus delivery (Sanworks), peristaltic pump for reward delivery, and an IR LED and photodetector as a beam break (Sanworks). Behavioral control software to implement the task and control individual boxes was written in MATLAB. Boxes were controlled through a Teensy-based microcontroller system (Bpod Sanworks). A custom written python application was used to control multiple Bpod instances from a single control computer, requiring an edited version of the Bpod MATLAB software library (edited Bpod library: https://github.com/RatAcad/Bpod_Gen2; Custom python application: https://github.com/RatAcad/BpodAcademy). Further details about the software implementation can be found in the respective code repositories. The floor of the chambers contained bedding.

### Rodent daily training

Mice were housed in groups of 2-4 mice per cage whereas rats were housed in pairs in an animal facility. Both were moved to the training room in the laboratory each morning for training. Mice ran for 1 hour shifts and rats ran for 2 hour shifts, 5 days per week in the late morning (10am-noon) or early afternoon (12pm-3pm). Rat feeding was conducted in the late afternoon after behavioral training (1-4pm).

### Rodent training pipeline

Rats and mice were rewarded with 10% sucrose at all stages of training and testing. Training took 1-4 weeks depending on the rodent and progressed through 3 stages. In the first stage (1-3 days) rodents were rewarded for inserting their nose into a side port with an illuminated LED (400 trials). In the second stage rodents received reward for inserting their nose into the center port and then the side port with an illuminated LED (400 trials). In the third stage, they received a reward for inserting their nose into the center port and then the side port with a flashing LED. Once rodents reached criterion 90 −100% correct over 400 trials, the probability of flashes on the incorrect side increased in the following way 100:0 -> 90:10 -> 80:20. Progression through this third stage took 1-5 days per condition. After completion of all stages and conditions rodents performed the task with a flash probability of 80:20.

### Rodent behavioral task

Rodents were free to move within their cage during the task. At the start of a trial, the light in the center port turned on indicating that the rodent could initiate the trial at its leisure. Once the rodent nose poked in the center port, a single flash would occur simultaneously in both the left and right ports. After this one light flash on both sides, the rodent would see a series of light flashes according to a Bernoulli process – every 100 ms, the rodent would see a flash on either the right or the left side with a probability of 80:20 that the flash would occur on the correct side vs. the incorrect side. The correct side was drawn randomly on each trial. To record a response, the rodent nose poked in either the left or right port and the series of light flashes would terminate as soon as this decision was recorded. If the rodent responded correctly, the light in the correct side port turned on for 3 s and a 30uL-rat or 5uL-mice sucrose water reward was delivered immediately. If the rodent responded incorrectly, all lights turned off for a 5 s time-out and no reward was delivered, and new trials would start after a 2s delay. Trials had an external response deadline of 8 seconds, after which it would be counted as an omission.

### Human participants

All experiments and procedures were performed in accordance with protocols approved by the Boston University Institutional Review Board. Players consisted of 18 adolescent male and females, ages 11-17, who had previous experience with different online non-verbal evidence accumulation games (e.g. Do et al. 2023). Players with a history of seizures were excluded. Players were recruited from Boston University and SPARK, had at least one sibling with a clinical diagnosis of autism, however, players themselves did not have an autism diagnosis.

### Human video-game and experimental procedure

The video game was developed using a JavaScript application called PixiJS, and run online using the cloud platform Heroku; data was saved online using the no-SQL database application MongoDB, in JSON format. Demographic and survey data were collected separately using Boston University’s REDCap platform. Each online game session was supervised by a member of the lab, trained in human behavioral testing procedures. This experimenter was present on video call (zoom) and provided guidance on the game setup, game rule was not disclosed; sessions were recorded. During the game, players destroyed asteroids. The first 10 trials served as learning trials, where participants needed to achieve above 90% accuracy to be included in the study. The remaining trials were test trials (80:20 generative flash probability). In each trial, participants selected the asteroid they wanted to destroy. Subsequently, a series of alien-shaped stimuli were presented according to a Poisson process, with an 80% probability of appearing on the correct side and a 20% probability of appearing on the incorrect side, or not at all, every 100 ms. The correct side was randomly determined for each trial. Participants responded by selecting the alien on the correct side, concluding the stimuli presentation. A correct response resulted in the destruction of the asteroid, awarded points, and a wait time of approximately 1 second before the next trial. An incorrect response led to the asteroid remaining intact, no points awarded, and a wait time of approximately 3 seconds before the next trial. If participants failed to respond within 8 seconds, the trial was counted as an omission.

### Synchronization of the free response pulse-based evidence accumulation task across species

We developed a cross-species free-response evidence accumulation task originally developed for rodents (Kane et al. 2023). In this task two competing streams of visual pulses were presented on the left and right sides of the subject’s visual field. Subjects were rewarded to select the side with a highly likelihood of pulsing to receive a reward. Pulses were generated using a Poisson process for humans and a Bernoulli process for rodents, with the pulse probability manipulated across task stages. The task maintained consistent stimulus statistics, including flash duration, flash rate, and generative probability, for both rodents and humans—specifically, a 10 Hz flash rate and an 80% flash probability on the rewarded side versus a 20% flash probability on the other side in the testing stage.

The task structure remained uniform across species, comprising four stages: initiation, free-response cue period, choice, and reward/time-out (Figure 1A). Human participants engaged in a single session (consisting of 200 trials) of the video game online under the supervision of a researcher via video call (Figure 1B). Multiple rodents performed the task in a 3-port chamber on a daily basis, with data transferred on the same day using Datajoint (Figure 1C).

### Data analyses

Data analysis was conducted using Python version 3.7.1 and R version 4.3.1. Multiple sessions were introduced per rodent, resulting in an average of 3605 ± 231 total trials per mouse and 4107 ± 549 total trials per rat. Chance performance of the task was determined by calculating the upper bound of the 99% binomial confidence interval for each species performing at chance, yielding an estimated chance performance of 52% for rodents and 60% for humans.

Accuracy was calculated by dividing the number of correct trials by the total number of trials and then multiplying the result by 100. Response time is defined as the average duration between the initiation of a stimulus and the subsequent response. Percentage bias was calculated by extracting the absolute difference between right and left trials and dividing by the total number of trials, the result was multiplied by 100. The average accuracy, response time, and reward rate during a session and across sessions were computed using a rolling average of 5 minutes during the 60 (mouse) or 120 (rat) minutes of the session. Trials across the session were calculated by counting the trials per minute and averaging them across sessions and subjects (Supplementary Figure 8). Standard statistical tests (e.g., One-way ANOVAs, Pearson and Spearman correlations, multiple comparisons and t-students) were conducted using the pingouin package (Vallat, 2018).

To model the accuracy versus response time relationship shown in Figure 2F, a generalized additive mixed model (GAMM) was utilized, employing the mgcv package (Wood et al., 2017). The data points on the plot represent the average accuracy of all trials at each response time, calculated using 0.2-second bins over a span of 3 seconds; only bins that presented more than 50 trials were selected. Pearson correlations were performed using the bins within the first 1.5 seconds for each species. All trials for each species were collectively fitted within the model framework and the data.

### Drift diffusion models

Drift diffusion models (DDMs) are widely utilized to characterize decision-making in two-choice paradigms. The DDM assumes that decision-makers accumulate noisy evidence for each choice option over time, ultimately committing to a response when a predefined decision threshold is reached. In our study, we employed three variants of DDMs: the classic DDM (Bogacz et al., 2006; Ratcliff, 1978; Ratcliff & McKoon, 2008), the collapsing boundary DDM (Bowman et al., 2012; Hawkins et al., 2015; Shevinsky & Reinagel, 2019; Voskuilen et al., 2016) and the urgency DDM (Ditterich, 2006; Hawkins et al., 2015). The classic DDM assumes that the amount of evidence required to reach a decision is fixed over time, and implements it with fixed decision boundaries (*a*). By convention, we assume the lower boundary to be 0, which sets a total boundary separation of *a* − 0 = *a*. In addition, it includes the following parameters:

1) The drift rate (*v*), which is the rate at which information is sampled; evidence (*x*) grows linearly with time (*t*): the change in evidence (*dx*), is updated by the drift rate (*v*), and a gaussian noise (*W*) scaled with a constant (*c*): *dx* = *v_i_* * *dt* + *cdW* Here, *i* represents a trial.
2) The non-decision time (*ndt*), which is the time taken up by processes not-related to evidence accumulation.
3) The starting point bias (*z*), which determines the location on the distance between the two boundaries from which the evidence accumulation process starts, it reflects a bias to choose one option over the other one. *z* is relative to *a*.

Thus, evidence is accumulated over time using the drift rate embedded with noise. Evidence is accumulated until the decision threshold is reached. We fit this model to the choice-RT data of individuals from each species. For each model we also assumed that drift rate, non-decision time and starting point bias can vary from trial-to-trial following normal, half-normal and uniform distributions, respectively:

Drift rate: *v_i_ ∼ N(v, theta_v_);*

*Starting point: x*0*_i_ ∼ Unif (x*0 - *theta_x_*0, *x*0 + *theta_x_*0);

*Non-decision time: ndt_i_ ∼ Unif (ndt* - *theta_n_dt*, *ndt + theta_n_dt*).

Unlike the classic DDM, both collapsing boundary model and the urgency model assume that the amount of evidence required to reach a decision is not fixed over time. The collapsing boundary model included three additional parameters— 1) initial boundary separation (*a*_0_), 2) the semi-saturation constant (*t*_0.5_), or the time when the boundary had collapsed by half, and 3) the total amount of collapse (*k*). The urgency model also included three additional parameters— 1) the slope of a (linear) urgency signal (*u*_*slope*_), 2) total amount of urgency (*u*_*mag*_), and 3) the onset delay of urgency (*τ*).

The basic idea behind fitting a DDM to experimental data is to find a set of parameters which can simulate trials with the same accuracy and RT distributions as the experimental data. This is done iteratively, by perturbing the parameters, starting from an initial set of values, and comparing the fit between the simulated and experimental trials. Here, we used the *χ*^2^ method to compare between the RT histograms of the simulated- and experimental data, because the method is relatively fast and less prone to outlier RTs (Ratcliff & Tuerlinckx, 2002). All models were fitted using a custom R package (rddm; https://github.com/gkane26/rddm).

### Simulation analyses

For all simulations with the perfect accumulator, we included 10,000 trials for each choice of RT; RT typically varied between 100 ms and 4 s, with an increment of 200 ms. For each choice of RT, we calculated the accuracy and/or reward rate of the accumulator, considering the 10,000 trials. Consistent with the current task, each trial had a generative flash probability (*p*) of 80%. For each trial, the correct side (left or right) was pre-determined with a coin flip. Individual flashes were generated with a Bernoulli process for the simulations for rodents and a Poisson process for the simulations for humans. Sequence of flashes continued only up to the choice of RT. The perfect accumulator always chose the side with greater total number of flashes. If the trial ended with the same flash counts on each side, the perfect accumulator randomly picked a side. We also considered a noisy accumulator, which was informed by the accuracy of each species at specific RTs; this was obtained from the accuracy predictions of the GAMM (Supplementary Figure 7).

For simulations of the reward-rate, in addition to RT, we considered species-specific choices of inter-trial intervals (ITI). As mentioned in the results, ITI was the interval between the response and the next stimulus onset; it included the time to initiate a trial and the time spent in post-response events, such as rewards, timeouts etc. The average ITI was different across species, there was also a difference in the average ITI for rewarded and unrewarded trials. These differences were accounted for in the simulations. In the current experiment, each correct response produced the same amount of rewards, assuming each reward to be of 1 *unit*, a total accuracy of *N* would produce a total of *N* rewards. For each choice of RT, reward-rate was calculated for the 10,000 trials as:

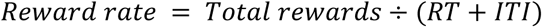

The optimal RT was the RT corresponding to the peak reward-rate in the RT versus reward-rate plot. Note that these simulations assumed that RT included the decision- or evidence-accumulation process only, whereas other models have also incorporated periods of non-accumulation (Simen et al., 2009).

### GLM and GLM-HMM

The generalized linear model (GLM) was utilized to elucidate choice probability, specifically the tendency to select the right option over the left option. This model incorporated three main predictors as a linear function:

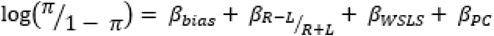

1. Pulse or flash difference: Calculated as the disparity between flashes on the two sides, normalized by the total number of flashes on both sides.
2. Win-stay or lose-switch strategy: Reflecting the tendency to stick with a chosen side after a win or switch sides after a loss.
3. Previous choice strategy: Indicating the inclination to follow the previous choice made.

The intercept term in the model represented the inherent bias towards one side over the other. Notably, this analysis was restricted to data from the 80% versus 20% flash stage.

To compare model fit across species with varying trial numbers, we employed the log-likelihood per trial in bits, as outlined by Ashwood et al. (2022). This is calculated as follows:

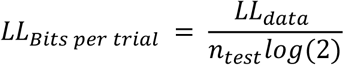

where *LL*_*data*_ is the log-likelihood of the entire data per species. *n*_*test*_ is the number of trials in the data. Predictive accuracy, following the methodology of Ashwood et al. (2022), was also utilized to gauge the goodness of fit.

Afterward, we compared these fits with a 2-state classic lapse model, which posits that subjects transition between two latent states. In one state, termed the engaged state, decisions are influenced by the stimuli, leading to dependent probabilities associated with the left and right stimuli. In the other state, referred to as the disengaged or lapse stage, decisions are entirely independent of the stimuli. In this scenario, subjects essentially flip a biased coin on each trial, with a fixed probability (γr+γl), and subsequently adopt one of two strategies based on the outcome: a strategy contingent on the stimulus or a strategy that disregards the stimulus entirely.

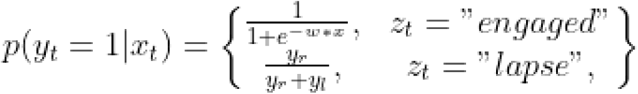

Furthermore, to elucidate potential transitions through different task stages, we applied a modeling framework incorporating hidden Markov models (HMMs). This framework, known as GLM-HMM, assumes distinct latent states that animals may inhabit during task performance, each potentially influencing decision strategies differently. This model comprises multiple independent GLMs, each corresponding to a latent state and characterizing choice outcomes as a Bernoulli process. Transition between states is governed by transition probabilities, and weight vectors within each state capture the relative influence of decision strategies. We fitted the GLM-HMM to cross-species data, varying the number of latent states and employing hyperparameters σ = 2 and α = 2 to govern the prior. A detailed description of the model is presented in Ashwood et al., (2022),

## Abbreviations

DDM: drift diffusion model
GAMM: generalized additive mixture model
GLM: generalized linear model
GLM-HMM: generalized linear model-hidden markov model
ITI: inter-trial interval
RT: response time

## Supplementary information

**Supplementary Figure 1:**
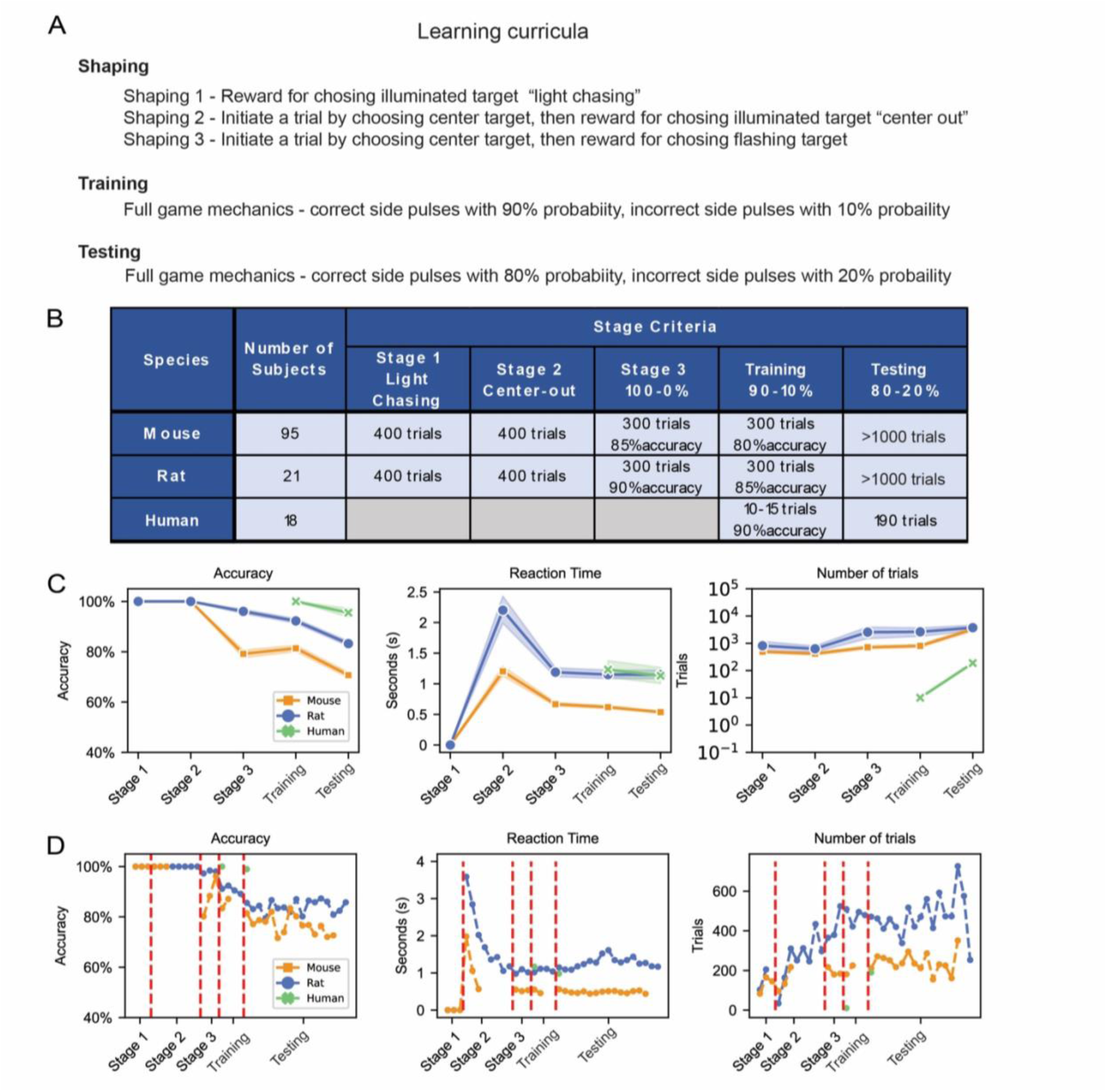
Learning curricula in the cross-species task. A) Experiments occurred in multiple phases: Shaping (rodents only), Training and Testing. Prior to training, rodents transitioned through three shaping stages to familiarize them with the operant chamber. In shaping stage 1 they received reward for inserting their noise in an illuminated noise poke. In shaping stage 2 they received reward for first poking in the center port to initiate a trial and then in the illuminated side port with the light. In shaping stage 3 they received reward for first poking in the center port to initiate a trial and then in the flashing side port. During the first stage of training both humans and rodents performed a simple version of the task in which the probability of cue presentation was high on the correct side (90% per time point) and low on the incorrect side (10% per time point). After completion of training, subjects transitioned to the testing phase in which the probability of cue presentation was 80% on the correct side, and 20% on the incorrect side. B) Table showing the number of subjects and training criteria for each species across the learning curricula. C) Evolution of averaged accuracy, response time and number of trials across learning in mice, rats and humans. D) Example mouse, rat and human evolution across stages and sessions. The red dashed lines represent the transitions across stages.

**Supplementary Figure 2:**
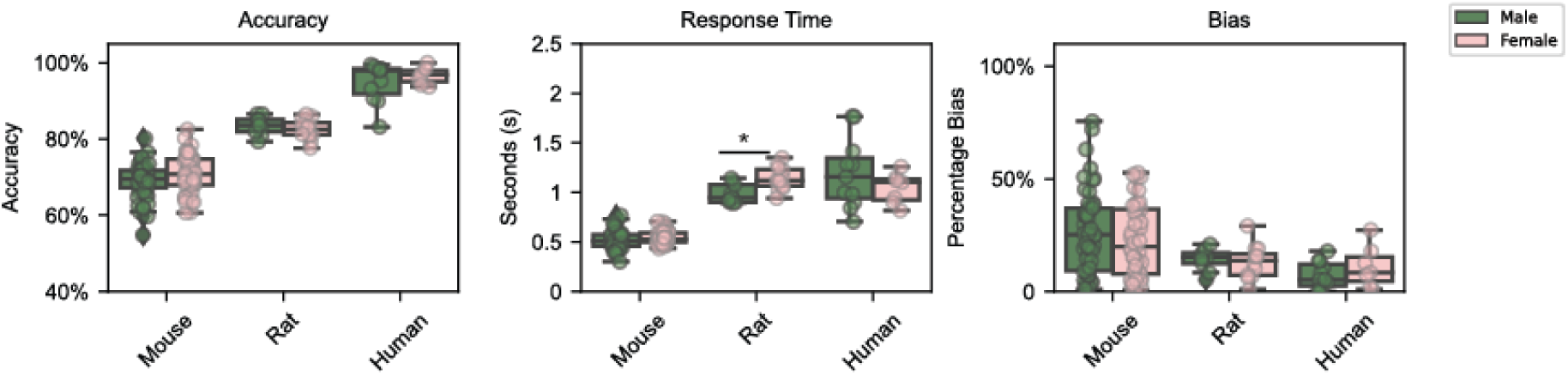
Comparison of accuracy, response time and bias across species, separately for female and male subjects. Notations: * = p<0.05, ** = p<0.01, *** = p<0.005, **** = p<0.001.

**Supplementary Figure 3:**
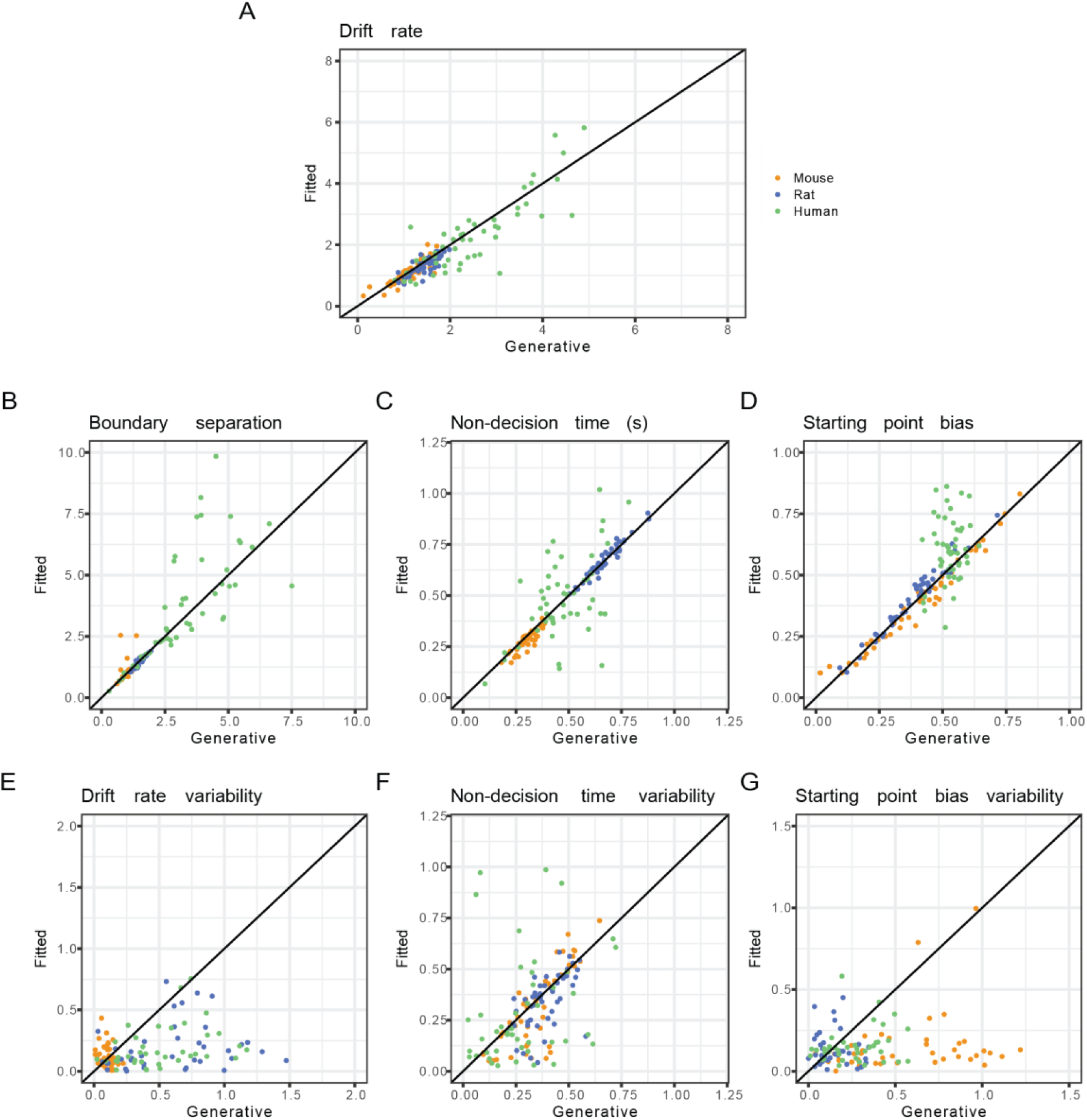
Parameter recovery for the DDM. Based on the parameters observed with the empirical data, we simulated 50 datasets per species with 600 trials each. After fitting the DDM to each of these simulated datasets, we compared the generative and fitted parameters.

**Supplementary Figure 4:**
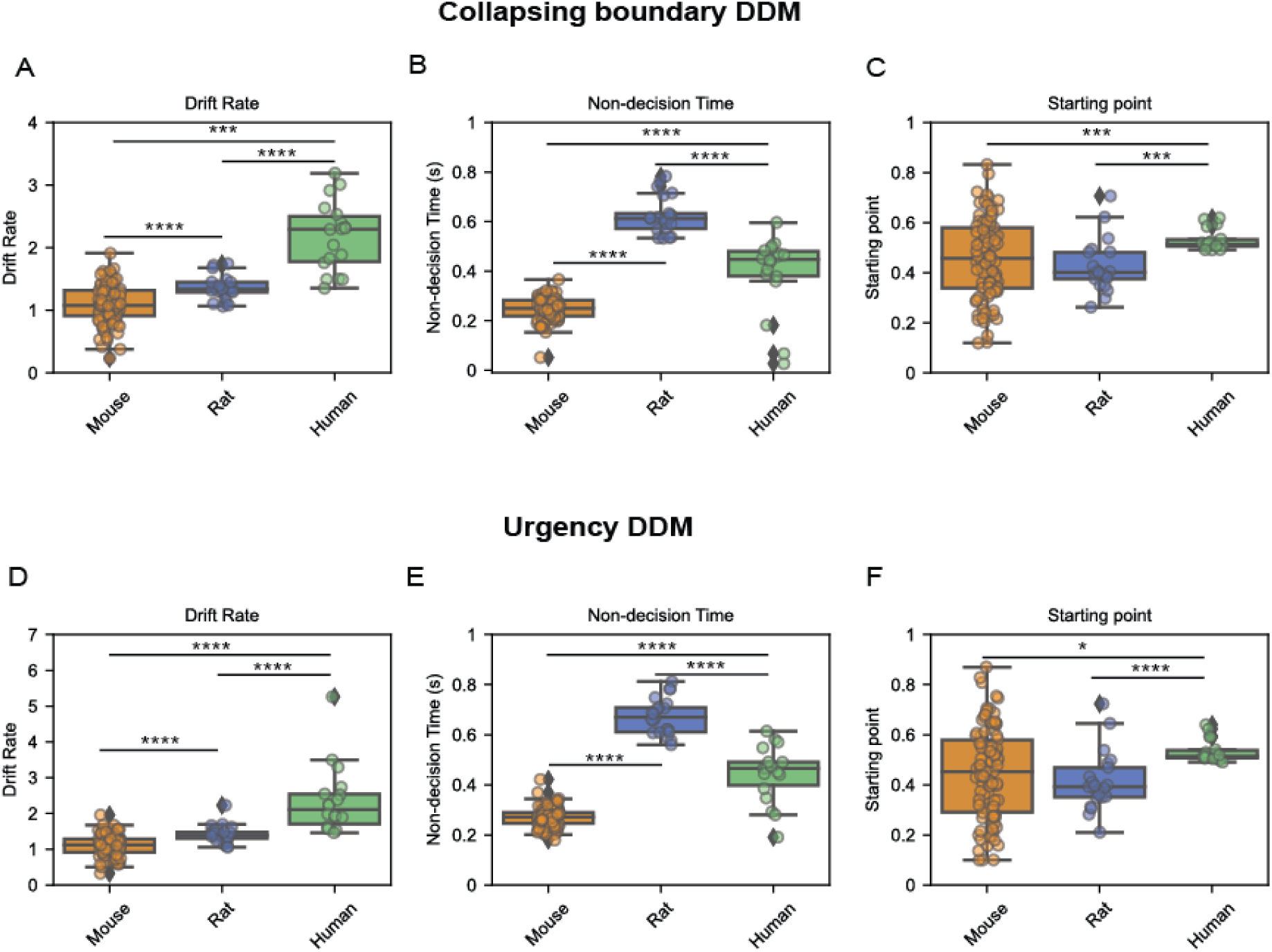
Comparison of drift rate, non-decision time and starting point bias across species, separately for the collapsing boundary DDM and the urgency DDM. Both models present similar parameter estimates and cross-species trends as the base DDM. Notations: * = p<0.05, ** = p<0.01, *** = p<0.005, **** = p<0.001.

**Supplementary Figure 5:**
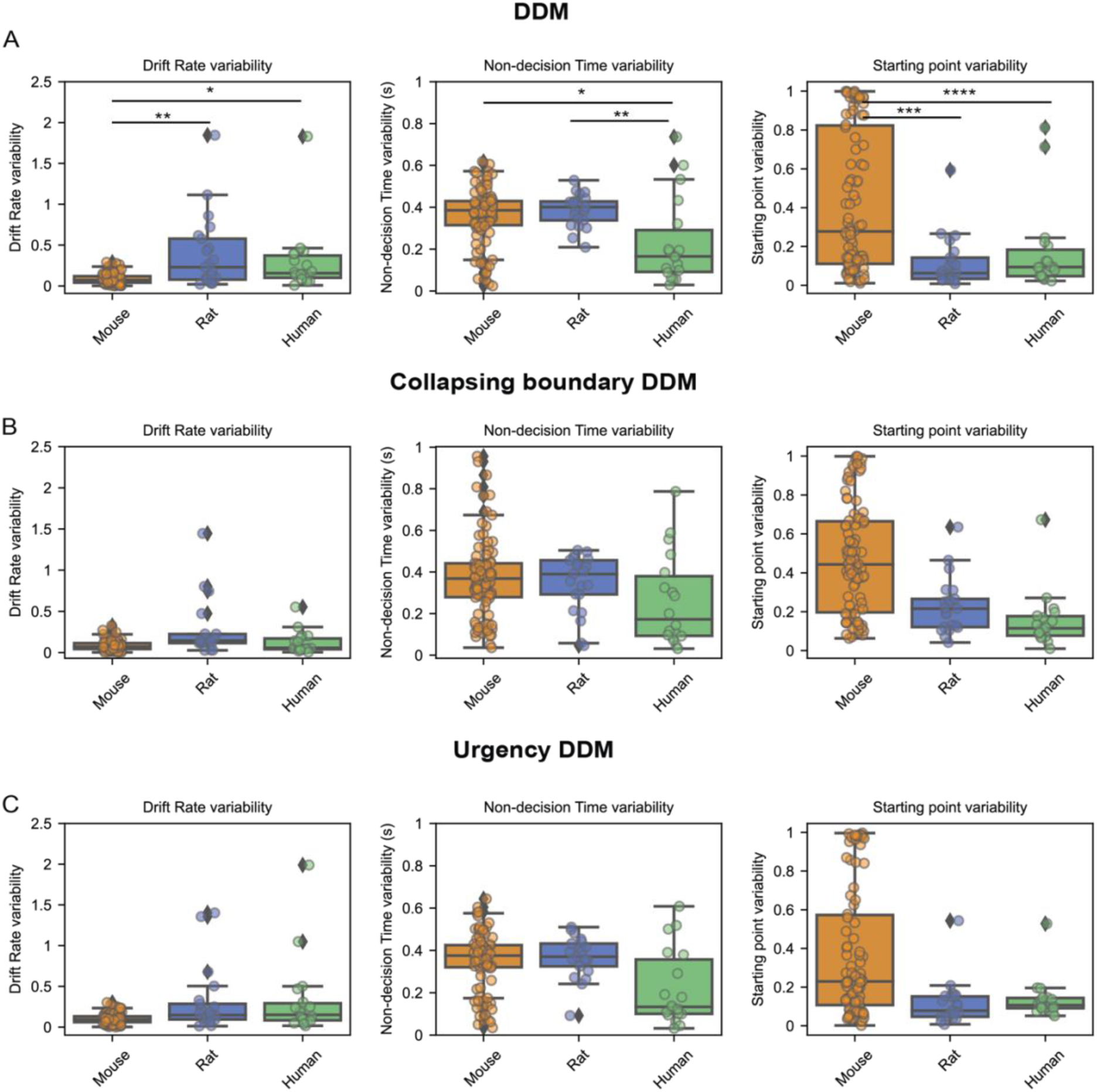
Comparison of DDM parameters that control the trial-to-trial variability of its main parameters, such as drift rate, non-decision time and starting point bias. We assumed variability of drift rate from trial to trial follows a normal distribution, that for non-decision time follows a half-normal distribution, and for starting point bias, trial-to-trial variability follows a uniform distribution. A) corresponds to the original DDM, B) corresponds to the collapsing boundary model, C) corresponds to the urgency DDM. Notations: * = p<0.05, ** = p<0.01, *** = p<0.005, **** = p<0.001.

**Supplementary Figure 6:**
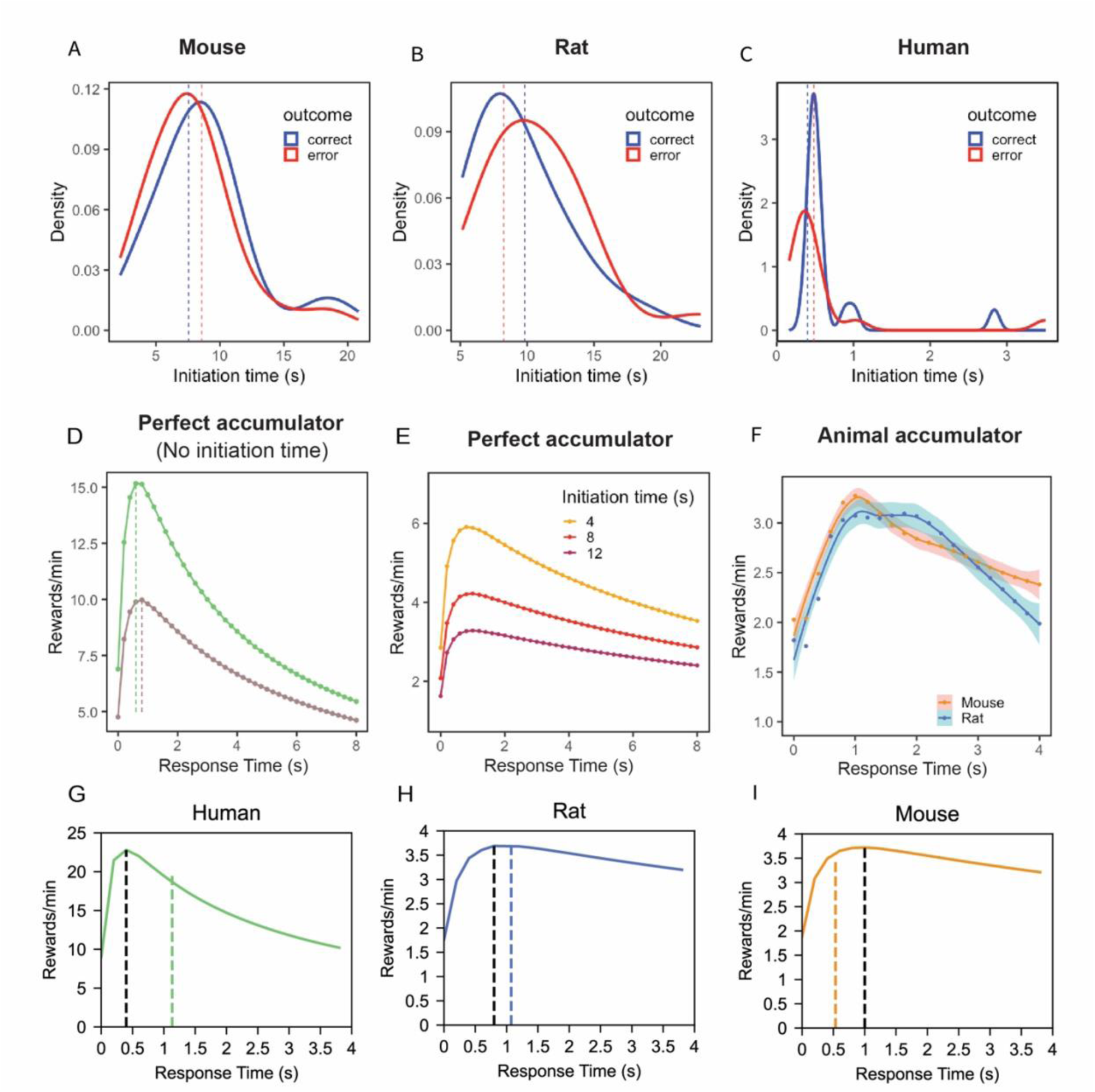
Differences in intertrial intervals across species and its effects on the reward rate simulations. A-C) Distributions of initiation times across all correct and erroneous responses, separately for mice, rats and humans. Initiation time was the period between the start of a trial and the center poke, after the center poke, flash stimuli were presented; for humans this was the time to choose an asteroid. D) Reward rate (rewards/min) as a function of different choices of RT for a perfect accumulator, simulated separately for humans (green line) and rodents (purple line). RT corresponding to the peak reward rate is noted. E) Reward rate of a perfect accumulator taking into account different initiation times. F) Reward rate as a function of different choices of RT for noisy (animal) accumulators. Separate simulations were run to account for average trial initiation times observed in rats and in mice, for correct and error responses. In the simulations, flash ratio was kept at 80% vs 20%, and flash rate was 10 Hz, same as that for the actual experiments. In addition to initiation time, choice accuracy for at different RT levels was predicted by the GAMM, previously fit to animal choice/RT data (see Figure 2F). G-I) Reward rate as a function of different choices of RT for humans (G), mice (H) and rats (I) based on a perfect accumulator, incorporating the average trial initiation time observed in each species, for correct and error responses. The black dashed line represents the response time where the reward rate of the accumulator peaks and the colored line is the average response time of each species.

**Supplementary Figure 7:**
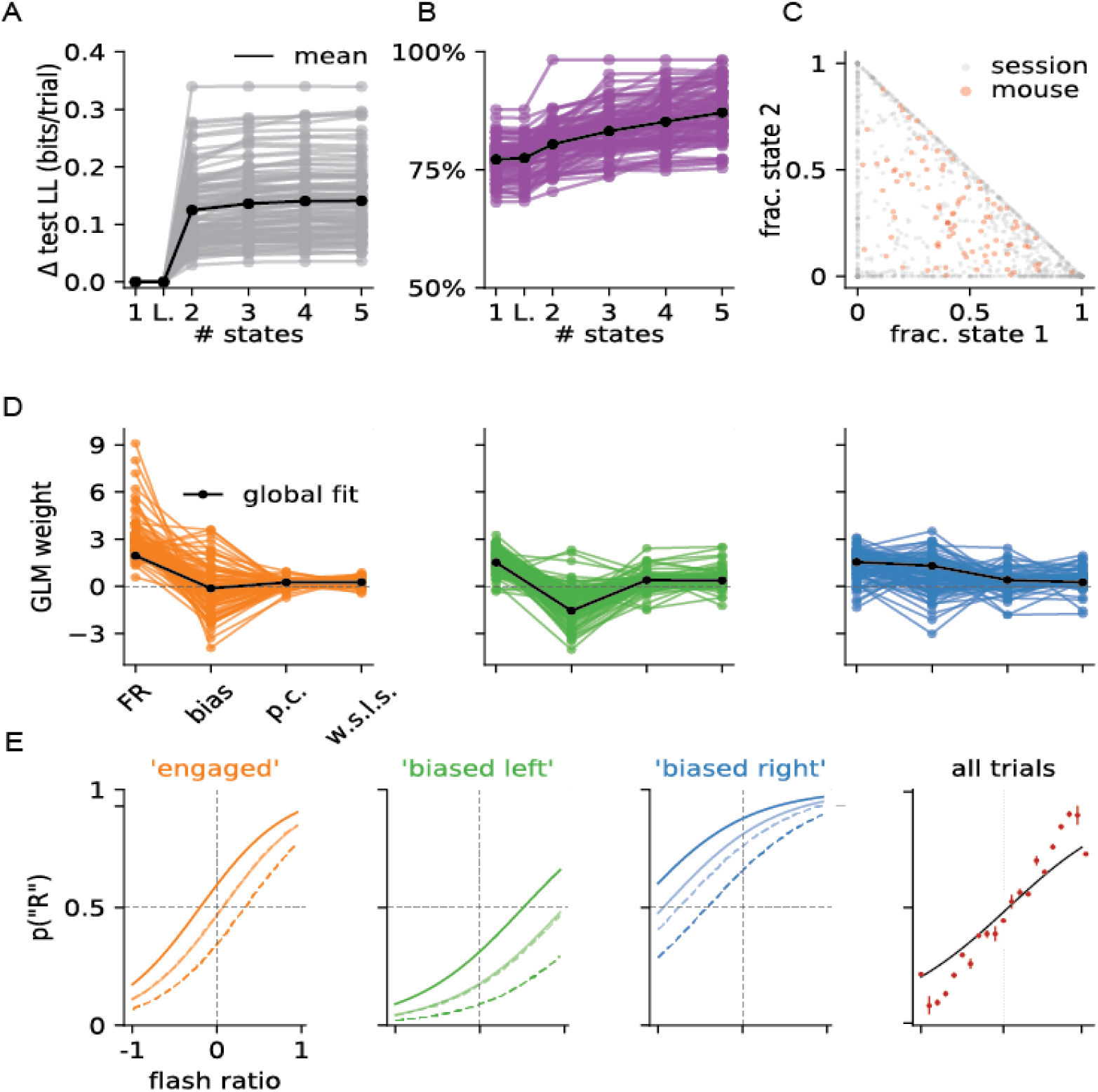
Mouse data is better fit with a hidden Markov model with 3 latent states. A) Change in test log-likelihood as a function of the number of states included in the GLM-HMM, relative to a (one-state) GLM, modeled separately for each mouse. The classic lapse model, a restricted form of the two-state model, is labeled as ‘L’. Each trace represents a single mouse and the solid black line indicates the mean across animals. B) Change in predictive accuracy relative to a one-state GLM for each mouse, indicating the percentage improvement in predicting choice. C) Gray dots correspond to individual sessions across all mice, indicating the fraction of trials spent in state 1 (engaged) and state 2 (biased left). Points at the vertices (1,0), (0,1) or (0,0) indicate sessions with no state changes, whereas points along the sides of the triangle indicate sessions that involve only two of the three states. Red dots correspond to the same fractional occupancies for each of the mice, revealing that the engaged state predominated but that all mice spent time in all three states. D) Inferred GLM weights for each mouse, for each of the three states in the three-state model. The solid black curve represents a global fit using pooled data from all mice. E) Psychometric curve for each state, conditioned on previous reward (solid line if right and dashed line if left) and previous choice (darker color if rewarded and dashed line if not rewarded) and all the states combined.

**Supplementary Figure 8:**
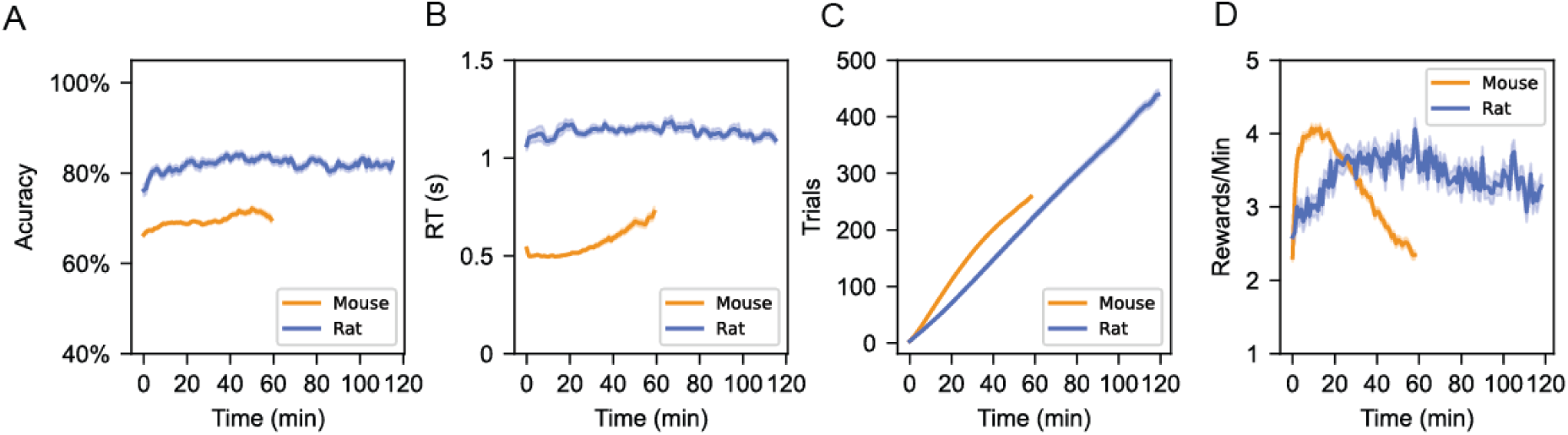
Rats exhibited stability throughout the entire session, whereas mice displayed greater variability. A-D) Averaged accuracy (A), response time (B), number of trials (C), and reward rate (D) across a session. It is important to note that mouse sessions lasted for 60 minutes and rat sessions for 120 minutes.

